# Understanding the genetics of important traits in quality protein maize (*Zea mays* L.) by Line × Tester analysis

**DOI:** 10.1101/2020.05.26.118125

**Authors:** Baudh Bharti, R.B. Dubey, Arun Kumar, Lalit Pal, Prashant Kaushik

## Abstract

Maize (*Zea mays* L.) is among the top ten most valuable crops and livestock products. Maize demand and production are continuously rising. The nutritionally rich quality proteins maize (QPM) has almost two times as much lysine as well as tryptophan, amino acids which then the conventional maize varieties. In this study, genetics of important traits in quality protein maize were determined using a Line by Tester matting design. The 45 hybrids were obtained via Line by Tester crossing of 15 lines and 3 testers. The parents and their hybrids were evaluated under two environments. A significant amount of variation was recorded for most of the traits studied. Parental genotypes L2, L6, T2, L8, L7 and L14, exhibited negative and considerable general combining ability (GCA) effects for days to flowering and maturity. L7 × T2, L13 × T3, L10 × T1, L1 × T1, L9 × T2, L5 × T1, L12 × T2, L11 × T3, L10 × T3, L14 × T1 and L6 × T1 were identified with suitable yield and component traits. Crosses L5 × T1, L3 × T1, L12 × T2, L7 × T3, L1 × T2 and L13 × T1 were identified for quality traits. Heterosis over the mid parent crosses L13 × T1 showed highest negative and significant followed by L6 × T1, L12 × T2, L14 × T2 and L2 × T1 for days to 50 per cent tasselling. A significant correlation for the lysine content was determined between F1 mean and specific combining ability (SCA) effects. Overall, this work provides the useful insights into the genetics of important agronomical and biochemical traits of quality protein maize.

## Introduction

Maize (*Zea mays* L.) is one of the most valuable crops in the world, and the demand of maize production is increasing in recent years since it has been extensively used to feed animals and as a new source of bioenergy. It is used worldwide for feed, food. Also, it serves as the source of the basic raw material of many industries *viz.,* oil, starch, protein, food, alcoholic beverages, sweeteners, cosmetics and biofuels etc. It is the principal field crop of the country. About 59% of the total production is used as feed, while the remaining is used as industrial raw material (17%), food (10%), export (10%) in other purposes (4%) [1,2]. Hence, genetic manipulation for improved nutritional value, particularly protein quality, was considered as a noble goal. This effort was stimulated by the 1963 discovery of mutant maize called as opaque-2 gene. The lysine content in the grains that are whole of quality protein maize (QPM) ranges from 0.33 to 0.54 per cent, with generally 0.38 as average and the tryptophan content is 0.08 per cent, which is 6.6 per cent higher than normal maize [3–5]. In maize, for example, the exploitation of heterosis or hybrid vigour contributes close to 40 % of grain yield boost in China [6]. There are two approaches in hybrid breeding; one of them is developing inbred lines, and then another one is selecting adequate parent inbred lines to mix elite hybrids [7].

Maize plays a crucial role in human and animal nutrition. In this direction, the breeding for enhanced protein quality in maize started in the mid-1960s with the discovery of mutants, like opaque 2, which produce enhanced levels of tryptophan and lysine, the 2 amino acids generally lacking in maize endosperm proteins. Nevertheless, undesirable pleiotropic consequences imposed severe constraints for effective exploitation of the mutants. The maize carrying opaque 2 gene in the homozygous state is actually described as opaque 2, and the tough modified endosperm opaque 2 maize with vitreous kernels is generally known as Quality Protein Maize (QPM). Composition of maize endosperm protein in addition to result in a two-fold rise in the amounts of tryptophan and lysine in comparison with the regular genotypes. Besides, other amino acids such as histidine, arginine, aspartic acid and glycine show an increase. At the same time, a decline is observed for some amino acids such as glutamic acid, alanine and leucine. The reduction in leucine is actually seen as especially appealing as it can make leucine isoleucine ratio much more healthy, which subsequently helps to liberate much more tryptophan for niacin biosynthesis and hence, helps to fight pellagra [8]. Inbreeding practice, it is well known that yield performance of hybrids could not be predicted by the performance of their parents per se [9].

On the contrary, it is primarily controlled by the potential to create superior hybrids or combining ability of the parental lines. Therefore, it is essential to select inbreds with high combining ability in hybrid breeding. Studies have suggested that the application of hybrids, especially single cross hybrids, leads to dramatically increasing of the yield of crops [10]. The combining ability partitioned into general combining ability (GCA), and specific combining ability (SCA), provides useful information regarding the decision of selection or hybrid development [11]. GCA of an inbred type is calculated as the typical functionality of all hybrids with that inbred type as a single parent, as well as SCA of certain combinations or maybe crosses is measured by the deviation of the hybrid performance from the GCA of the parents [12]. The current study was performed for estimating GCA, SCA, heterosis and the underlying gene effects for grain yield as well as quality traits in QPM maize hybrids over environments produced by the line × tester mating design [13].

## Methods and Materials

### Plant Material, Experimental Sites and Soil Properties

The experimental material was generated by making crosses between 15 inbred lines and 3 testers in line × tester mating design. Parents and their 45 hybrids were evaluated during Kharif of 2014-15 across locations. Three environments were created by two locations and date of sowing *viz*., E1 (timely sowing, Kharif 2014 at Instructional farm Rajasthan college of Agriculture, Udaipur), E2 (timely sowing, Kharif 2014 at ARSS, Vallabh Nagar, Maharana Partap University of Agriculture Science and Technology, Udaipur). The information pedigree of parents, checks and soil properties of these experimental sites are given in Supplementary Table-1. Geographically Udaipur is situated at 24° - 35’ North latitude and longitude 73° 42’ East at an elevation of 582.17 meters above mean sea level and metrological data were given in Supplementary Table-2.

### Experimental Design

The parent and their hybrids were planted under each environment in randomized block design with three replications in a single row plot of four-meter length, maintaining crop geometry of 60 × 25 cm. Randomization of genotypes was done by Crop Stat v7.2 software. All the recommended agronomy inputs and practices were applied to the crop during the season to raise the successful crop.

### Data Collection

Days to 50 % tasselling (counted as the number of days from sowing to when 50% of the tassel flowered), days to 50 per cent silking (counted as the number of days from sowing to when 50 % of the silk flowered), anthesis silking interval, days to 75 per cent brown husk (at the time when 75% of the plants reached physiological maturity) plant height (measured in cm from the base to tip of the tassel start to branch), ear height (measured in cm from ground level to node bearing the upper most ear), ear length (measured in cm from the bottom end to the tip of apical bud of the individual ear), ear girth (measured in cm from the thickness of the ear at the middle of the ear), number of grain rows per ear, the shelling percentage was calculated as ratio of grain weight to ear weight,100-grain weight (g), grain yield per plant (g), Harvest index was also calculated as ratio of grain yield and fresh biomass yield. Data was collected on 10 randomly selected and tagged crops for those characters except for days or weeks to fifty per cent tasselling, days or weeks to fifty per dollar silking, anthesis silking interval days or weeks to seventy-five per cent murky husk which was captured on a plot schedule.

### Quality Traits

Two random samples were drawn from the bulk harvest of ten self-pollinated plants under each replication.

*Oil content (%):* The Soxhlet’s ether extraction method defined elsewhere [14] was used for the estimation of oil content.

*Starch content (%):* The starch content was determined by the Anthrone Reagent method.

*Protein content (%):* Protein content of the seeds was estimated by using Micro Kjeldahl's method given by [15].

*Lysine content and Tryptophan content (%):* Calorimetric methods were used for the lysine and tryptophan content (%) content estimation as defines elsewhere [16, 17].

### Data Analysis

The combining ability effects for line × tester mating design was performed as per the method suggested by Kempthorne (1957) [13]. Heterosis and heterobeltiosis were calculated according to the method suggested by Shull (1908) [18], Fonseca and Patterson (1968) [19], heritability and genetic advance was estimated as per Lush (1949) [20] and Burton (1952) [21]. The analysis was done by Windostat version 9.2 and graphs were plotted by using Minitab v16.0 software. Correlation among values of the different traits investigated was performed based on Pearson’s product-moment correlation and cluster dendrogram as implemented in R (R Development Core Team2008). Descriptive statistics were done by SPSS v 16.0 software.

## Results

Combined estimation of variance revealed significant variations (p < 0.5) in the genetic material for traits studied in the present investigation (Table 1). Furthermore, significant differences were determined for genotype and its location interaction except for the traits days to 50 % tasselling, days to 75 % brown husk, plant height, ear girth, number of grain rows per ear, and 100-grain weight.Shelling percentage, harvest index, starch content and protein content suggesting the existence of varied genotypic performance across test locations. Line by location, tester by location, and line × tester by location line tester and line × tester main effects were significant for most traits. Highly significant line, tester and line × tester effects indicate the role of both additive and non-additive gene action in the inheritance of studied traits, in that order.

**Table 1.**
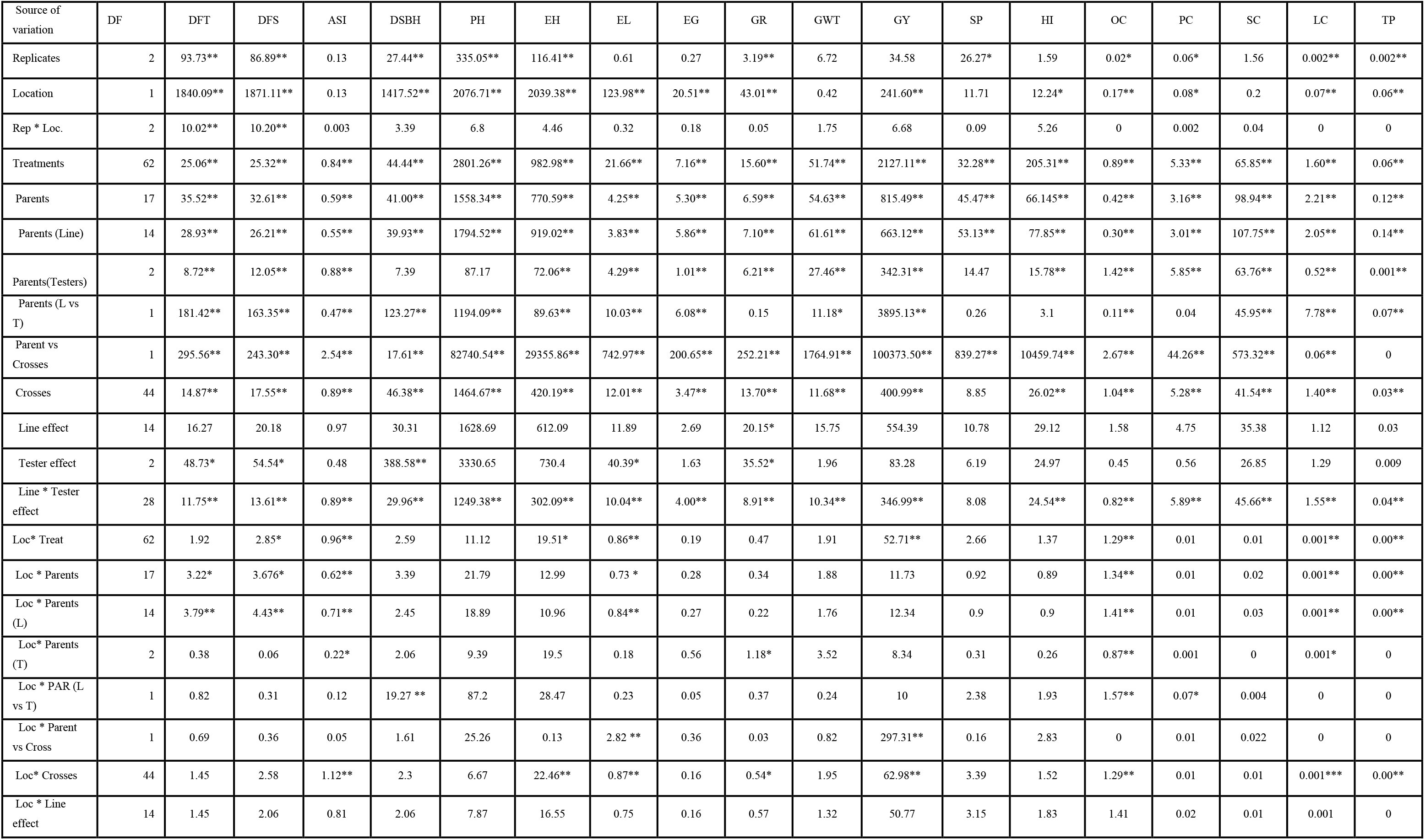
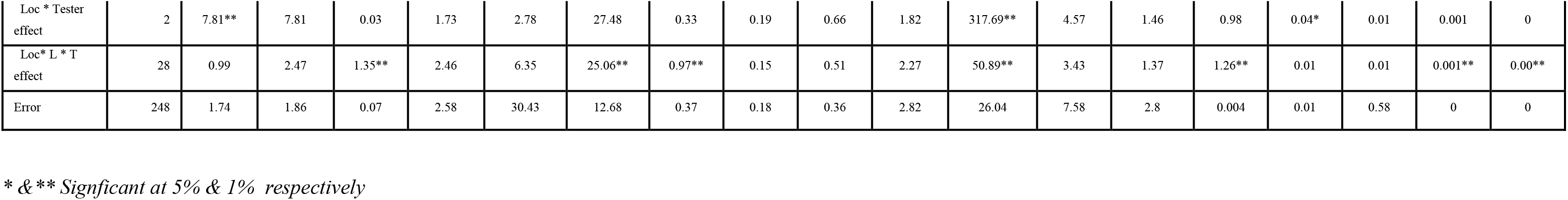
Analysis of variance for phenological, morphological grain yield and quality traits in QPM hybrids and their parents.

| Source of variation | DF | DFT | DFS | ASI | DSBH | PH | EH | EL | EG | GR | GWT | GY | SP | HI | OC | PC | SC | LC | TP |
| --- | --- | --- | --- | --- | --- | --- | --- | --- | --- | --- | --- | --- | --- | --- | --- | --- | --- | --- | --- |
| Replicates | 2 | 93.73** | 86.89** | 0.13 | 27.44** | 335.05** | 116.41** | 0.61 | 0.27 | 3.19** | 6.72 | 34.58 | 26.27* | 1.59 | 0.02* | 0.06* | 1.56 | 0.002** | 0.002** |
| Location | 1 | 1840.09** | 1871.11** | 0.13 | 1417.52** | 2076.71** | 2039.38** | 123.98** | 20.51** | 43.01** | 0.42 | 241.60** | 11.71 | 12.24* | 0.17** | 0.08* | 0.2 | 0.07** | 0.06** |
| Rep * Loc. | 2 | 10.02** | 10.20** | 0.003 | 3.39 | 6.8 | 4.46 | 0.32 | 0.18 | 0.05 | 1.75 | 6.68 | 0.09 | 5.26 | 0 | 0.002 | 0.04 | 0 | 0 |
| Treatments | 62 | 25.06** | 25.32** | 0.84** | 44.44** | 2801.26** | 982.98** | 21.66** | 7.16** | 15.60** | 51.74** | 2127.11** | 32.28** | 205.31** | 0.89** | 5.33** | 65.85** | 1.60** | 0.06** |
| Parents | 17 | 35.52** | 32.61** | 0.59** | 41.00** | 1558.34** | 770.59** | 4.25** | 5.30** | 6.59** | 54.63** | 815.49** | 45.47** | 66.145** | 0.42** | 3.16** | 98.94** | 2.21** | 0.12** |
| Parents (Line) | 14 | 28.93** | 26.21** | 0.55** | 39.93** | 1794.52** | 919.02** | 3.83** | 5.86** | 7.10** | 61.61** | 663.12** | 53.13** | 77.85** | 0.30** | 3.01** | 107.75** | 2.05** | 0.14** |
| Parents(Testers) | 2 | 8.72** | 12.05** | 0.88** | 7.39 | 87.17 | 72.06** | 4.29** | 1.01** | 6.21** | 27.46** | 342.31** | 14.47 | 15.78** | 1.42** | 5.85** | 63.76** | 0.52** | 0.001** |
| Parents (L vs T) | 1 | 181.42** | 163.35** | 0.47** | 123.27** | 1194.09** | 89.63** | 10.03** | 6.08** | 0.15 | 11.18* | 3895.13** | 0.26 | 3.1 | 0.11** | 0.04 | 45.95** | 7.78** | 0.07** |
| Parent vs Crosses | 1 | 295.56** | 243.30** | 2.54** | 17.61** | 82740.54** | 29355.86** | 742.97** | 200.65** | 252.21** | 1764.91** | 100373.50** | 839.27** | 10459.74** | 2.67** | 44.26** | 573.32** | 0.06** | 0 |
| Crosses | 44 | 14.87** | 17.55** | 0.89** | 46.38** | 1464.67** | 420.19** | 12.01** | 3.47** | 13.70** | 11.68** | 400.99** | 8.85 | 26.02** | 1.04** | 5.28** | 41.54** | 1.40** | 0.03** |
| Line effect | 14 | 16.27 | 20.18 | 0.97 | 30.31 | 1628.69 | 612.09 | 11.89 | 2.69 | 20.15* | 15.75 | 554.39 | 10.78 | 29.12 | 1.58 | 4.75 | 35.38 | 1.12 | 0.03 |
| Tester effect | 2 | 48.73* | 54.54* | 0.48 | 388.58** | 3330.65 | 730.4 | 40.39* | 1.63 | 35.52* | 1.96 | 83.28 | 6.19 | 24.97 | 0.45 | 0.56 | 26.85 | 1.29 | 0.009 |
| Line * Tester effect | 28 | 11.75** | 13.61** | 0.89** | 29.96** | 1249.38** | 302.09** | 10.04** | 4.00** | 8.91** | 10.34** | 346.99** | 8.08 | 24.54** | 0.82** | 5.89** | 45.66** | 1.55** | 0.04** |
| Loc* Treat | 62 | 1.92 | 2.85* | 0.96** | 2.59 | 11.12 | 19.51* | 0.86** | 0.19 | 0.47 | 1.91 | 52.71** | 2.66 | 1.37 | 1.29** | 0.01 | 0.01 | 0.001** | 0.00** |
| Loc * Parents | 17 | 3.22* | 3.676* | 0.62** | 3.39 | 21.79 | 12.99 | 0.73 * | 0.28 | 0.34 | 1.88 | 11.73 | 0.92 | 0.89 | 1.34** | 0.01 | 0.02 | 0.001** | 0.00** |
| Loc * Parents (L) | 14 | 3.79** | 4.43** | 0.71** | 2.45 | 18.89 | 10.96 | 0.84** | 0.27 | 0.22 | 1.76 | 12.34 | 0.9 | 0.9 | 1.41** | 0.01 | 0.03 | 0.001** | 0.00** |
| Loc* Parents (T) | 2 | 0.38 | 0.06 | 0.22* | 2.06 | 9.39 | 19.5 | 0.18 | 0.56 | 1.18* | 3.52 | 8.34 | 0.31 | 0.26 | 0.87** | 0.001 | 0 | 0.001* | 0 |
| Loc * PAR (L vs T) | 1 | 0.82 | 0.31 | 0.12 | 19.27 ** | 87.2 | 28.47 | 0.23 | 0.05 | 0.37 | 0.24 | 10 | 2.38 | 1.93 | 1.57** | 0.07* | 0.004 | 0 | 0 |
| Loc * Parent vs Cross | 1 | 0.69 | 0.36 | 0.05 | 1.61 | 25.26 | 0.13 | 2.82 ** | 0.36 | 0.03 | 0.82 | 297.31** | 0.16 | 2.83 | 0 | 0.01 | 0.022 | 0 | 0 |
| Loc* Crosses | 44 | 1.45 | 2.58 | 1.12** | 2.3 | 6.67 | 22.46** | 0.87** | 0.16 | 0.54* | 1.95 | 62.98** | 3.39 | 1.52 | 1.29** | 0.01 | 0.01 | 0.001*** | 0.00** |
| Loc * Line effect | 14 | 1.45 | 2.06 | 0.81 | 2.06 | 7.87 | 16.55 | 0.75 | 0.16 | 0.57 | 1.32 | 50.77 | 3.15 | 1.83 | 1.41 | 0.02 | 0.01 | 0.001 | 0 |
| Loc * Tester<br>effect | 2 | 7.81** | 7.81 | 0.03 | 1.73 | 2.78 | 27.48 | 0.33 | 0.19 | 0.66 | 1.82 | 317.69** | 4.57 | 1.46 | 0.98 | 0.04* | 0.01 | 0.001 | 0 |
| Loc* L * T<br>effect | 28 | 0.99 | 2.47 | 1.35** | 2.46 | 6.35 | 25.06** | 0.97** | 0.15 | 0.51 | 2.27 | 50.89** | 3.43 | 1.37 | 1.26** | 0.01 | 0.01 | 0.001** | 0.00** |
| Error | 248 | 1.74 | 1.86 | 0.07 | 2.58 | 30.43 | 12.68 | 0.37 | 0.18 | 0.36 | 2.82 | 26.04 | 7.58 | 2.8 | 0.004 | 0.01 | 0.58 | 0 | 0 |

\* & \*\* Significant at 5% & 1% respectively

### Performance of Maize parents and their Crosses

The mean performance revealed that the best-performing parents included L6 (70.53g/plant) T3 (64.63g/plant), T2 (57.45g/plant), L15 (50.57g/plant), T1(49.53g/plant) over the pooled location (Figure 1). The top five highest-yielding crosses were as follows: L7×T2 (92.12g/plant), L14 × T1 (91.33g/plant, L5×T1 (90.13g/plant), L6×T1 (90.00g/plant) and L12 × T2 (89.13/plant). Parents, L15 (27.60g/plant),L10(27.10g/plant), L6 (27.03g/plant), T1 (26.00g/plant) and L8 (24.30g/plant); and crosses L6 × T1 (31.87g/plant), L14 × T3 (30.40g/plant), L4 × T1 (29.63g/plant), L5 × T3 (29.43g/plant) and L2 × T2 (29.37g/plant) were the highest performing entries for 100-grain weight (Figure 1)‥ Among the parents, T1 (81.16 days) was the earliest to mature followed by T2 (81.333 days), L10 (82.00 days), L9 (82.67days) and L5 (82.83 days), while the crosses L14 × T2 (80.00 days), L8 × T2 (80.33days), L8 × T1 (81.17days), L5 × T2 (81.50 days) and L2 × T1 (81.67days) were the earliest to mature (Figure 1)‥ The cross L3 × T3 (93.17 days) was the latest maturing entry. L5 × T1 (5.53%) was showed highest for oil content, L1 × T2 (12.08%) for protein content, cross L7 × T3 for starch content (66.93%), L1 × T2 (2.37%) for lysine content (Figure 1). The crosses L8 × T1 (0.97) exhibited highest tryptophan content followed by L10 × T3 (0.69%), L5 × T1 (0.68%), L3 × T1(0.66%) and L4 × T2 (0.66%) (Figure 1).

**Figure 1.**
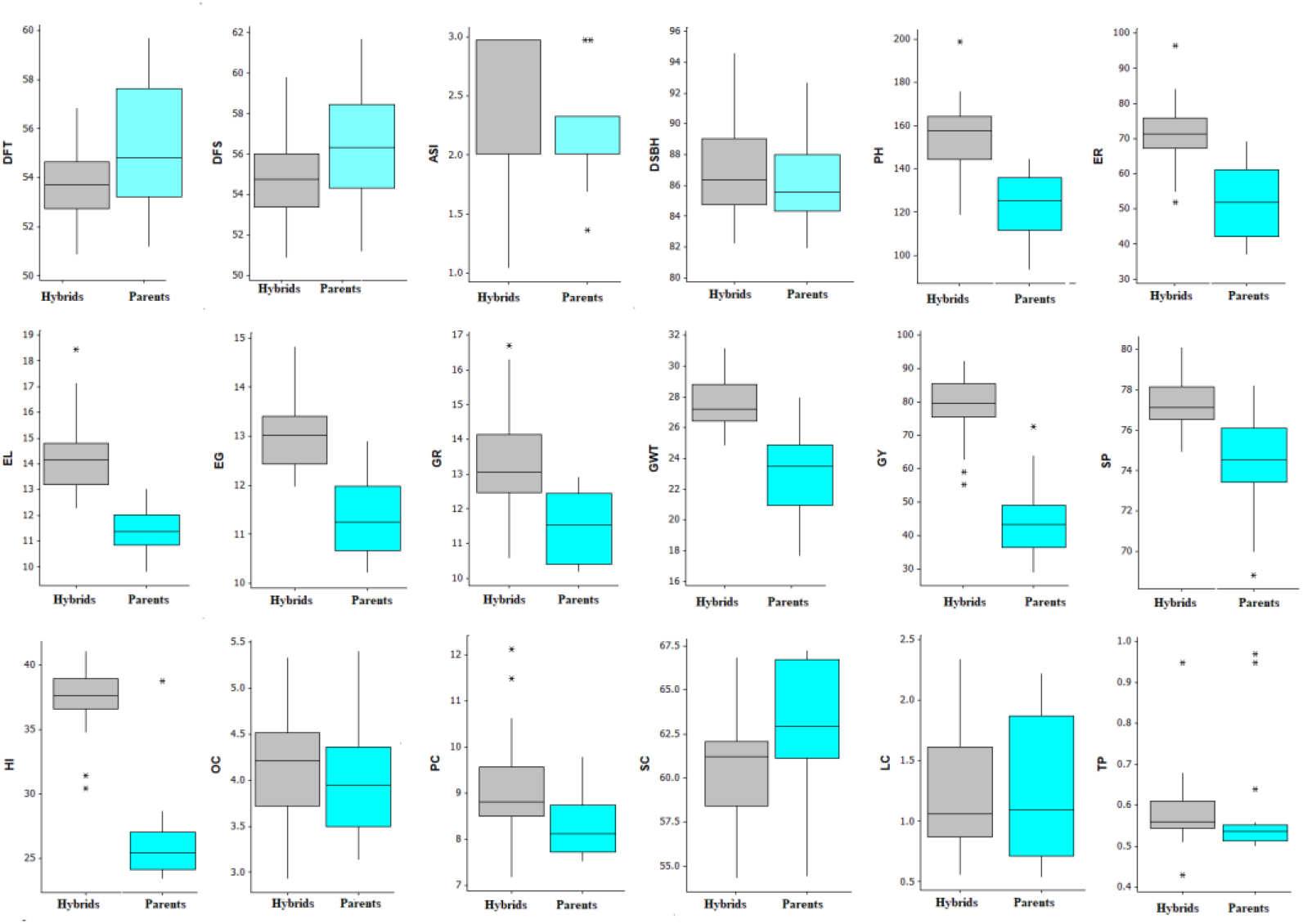
Box plot for grain yield and its contributing traits in QPM hybrids.

### Estimates of GCA and SCA Effects

Estimated values of GCA effects among parental maize genotypes, and SCA effects among crosses for yield, yield-related traits and quality traits are presented in Tables 2 and 3. Negative and highly significant GCA effects were recorded for L3, L2, L6, T2, L1 and L8 in a desirable direction for days to 50 per cent tasselling and parent L3, L2, L6, L1, L8, and T2 for days to 50 per cent silking reflecting their contribution to the breeding of early maturing varieties. Conversely, positive and highly significant GCA effects were observed for Days to 50 per cent tasselling for L11, L14, L5, L15, T3, L12 and L13 for the same trait, suggesting that these parents promoted late flowering in their crosses in an undesirable effect (Table 2). Parental genotypes L15, L7, L11, L12, L5, L9, T2 and L14 showed positive and significant GCA effects for starch content. For Lysine content positive and significant showed for parents L14, L4, L5, L12, L1, T2, T1, L8 and L11(Table 2). Positive and significant GCA effects were observed for tryptophan content among five parents, namely L8, L5, L14, L11 and T1 (Table 2).

**Table 2.** The general combining abilities (GCA) for phenological, morphological grain yield and quality traits in QPM hybrids over the location

| Parents | DFT | DFS | ASI | DSBH | PH | EH | EL | EG | GR | GWT | GY | SP | HI | OC | PC | SC | LC | TP |
| --- | --- | --- | --- | --- | --- | --- | --- | --- | --- | --- | --- | --- | --- | --- | --- | --- | --- | --- |
| L1 | -0.61* | -0.86** | -0.26** | -0.64 | 3.43* | 1.09 | 0.39** | -0.39** | 0.14 | -1.55** | -10.72** | -0.32 | -2.85** | 0.02 | 1.14** | -0.29 | 0.13** | -0.04** |
| L2 | -1.27** | -1.36** | -0.09 | -1.92** | -6.07** | -2.25** | -1.05** | 0.06 | -0.43** | 0.13 | 5.87** | -0.16 | 1.09* | -0.15** | -0.55** | -1.79** | -0.10** | -0.01* |
| L3 | -1.94** | -2.14** | -0.20** | 1.52** | 4.87** | 2.86** | -0.62** | -0.45** | 1.22** | -0.78* | 0.65 | -0.54 | -0.7 | -0.29** | -0.03 | -1.49** | -0.10** | 0.001 |
| L4 | -0.49 | -0.25 | 0.24** | 0.08 | 7.76** | 0.92 | 0.11 | 0.57** | -0.09 | 0.79* | 4.18* | 0.47 | 0.01 | -0.26** | -0.24** | -0.17 | 0.28** | -0.03** |
| L5 | 0.94** | 0.92** | -0.03 | 0.58 | 2.7 | 1.47 | -0.95** | -0.47** | -0.80** | -0.19 | 2.56* | -0.2 | 0.74 | 0.63** | 0.43** | 1.214** | 0.24** | 0.03** |
| L6 | -0.89* | -1.19 ** | -0.31** | -0.09 | 5.04** | -1.86* | -1.05** | 0.15 | -1.14** | 1.80** | 4.52** | -1.18 | 0.64 | -0.04* | -0.39** | -1.46** | -0.33** | -0.03** |
| L7 | -0.22 | -0.03 | 0.18* | -1.53** | -17.23** | -10.86** | -1.16** | 0.31* | -0.03 | -0.13 | -3.99** | 0.43 | -0.02 | 0.03 | 0.25** | 1.71** | -0.03** | -0.04** |
| L8 | -0.61* | -0.69* | -0.09 | -1.81** | -8.46** | -0.47 | 0.66** | -0.49** | 1.91** | 0.35 | 4.01** | -0.35 | 0.59 | 0.25** | 0.28** | 0.002 | 0.02** | 0.12** |
| L9 | 0.23 | 0.08 | -0.14* | -0.26 | -13.57** | -6.19** | -0.32* | 0.42** | -0.89** | -1.19** | -9.44** | 0.09 | -1.83** | -0.33** | -0.59** | 0.78** | -0.11** | -0.03** |
| L10 | 0.45 | 0.64* | 0.19** | 0.24 | -1.41 | -6.69** | 1.15** | -0.31* | -0.30* | 0.71 | -1.85 | -1.02 | -1.29** | 0.35** | -0.39** | -2.44** | -0.27** | 0.004 |
| L11 | 1.34** | 1.25** | -0.09 | 0.69 | -12.69** | -1.80* | 0.58** | 0.43** | 1.36** | -0.86* | -0.36 | 0.17 | 1.15** | 0.35** | 0.02 | 1.33** | 0.01** | 0.01** |
| L12 | 0.62* | 0.64* | 0.02 | 1.69** | 7.93** | -2.42** | 1.13** | -0.07 | 0.58** | -0.34 | 2.73* | 1.92** | 0.73 | -0.45** | -0.35** | 1.25** | 0.18** | -0.02** |
| L13 | 0.56* | 0.69* | 0.13* | 2.47** | 6.32** | 11.86** | 0.46* | -0.002 | -1.76** | -0.66 | -7.64** | -0.21 | -0.52 | -0.11** | -0.02 | -1.16** | -0.14** | -0.003 |
| L14 | 1.06** | 0.92** | -0.14* | -1.26** | 15.76** | 7.97** | 0.75** | 0.53** | 1.15** | 1.17** | 5.49** | 1.09 | 2.08** | -0.12** | 0.86** | 0.57** | 0.57** | 0.03** |
| L15 | 0.84** | 1.42** | 0.58** | 0.24 | 5.70** | 6.36** | -0.06 | -0.29** | -0.94** | 0.75 | 4.01** | -0.2 | 0.18 | 0.13** | -0.42** | 1.96** | -0.35** | 0.002 |
| T1 | -0.16 | -0.24 | -0.08** | -0.02 | 3.76** | -2.57** | -0.31** | 0.03 | -0.48** | 0.12 | 1.01 | 0.23 | -0.06 | -0.06** | 0.09** | -0.28** | 0.06** | 0.01** |
| T2 | -0.64** | -0.63** | 0.01 | -2.07** | -7.02** | -0.49 | -0.46** | -0.15** | -0.23** | -0.17 | -0.09 | -0.29 | -0.49** | 0.08** | -0.05** | 0.63** | 0.08** | -<br>0.004** |
| T3 | 0.80** | 0.87** | 0.07* | 2.09** | 3.26** | 3.06** | 0.77** | 0.12** | 0.71** | 0.04 | -0.91 | 0.08 | 0.55** | -0.02** | -0.04* | -0.35** | -0.14** | -0.01** |
| CD<br>0.05(Line) | 0.53 | 0.54 | 0.12 | 0.76 | 2.72 | 1.69 | 0.28 | 0.19 | 0.27 | 0.77 | 2.56 | 1.37 | 0.79 | 0.03 | 0.06 | 0.37 | 0.01 | 0.01 |
| 0.05 (Tester) | 0.24 | 0.24 | 0.05 | 0.34 | 1.22 | 0.76 | 0.13 | 0.09 | 0.12 | 0.34 | 1.14 | 0.61 | 0.36 | 0.02 | 0.03 | 0.17 | 0.002 | 0.002 |

**Table 3.**
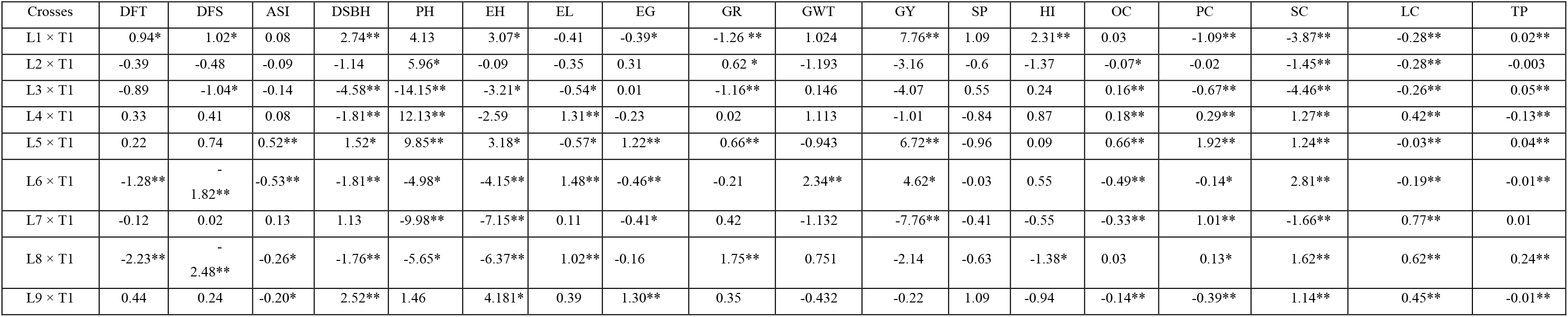
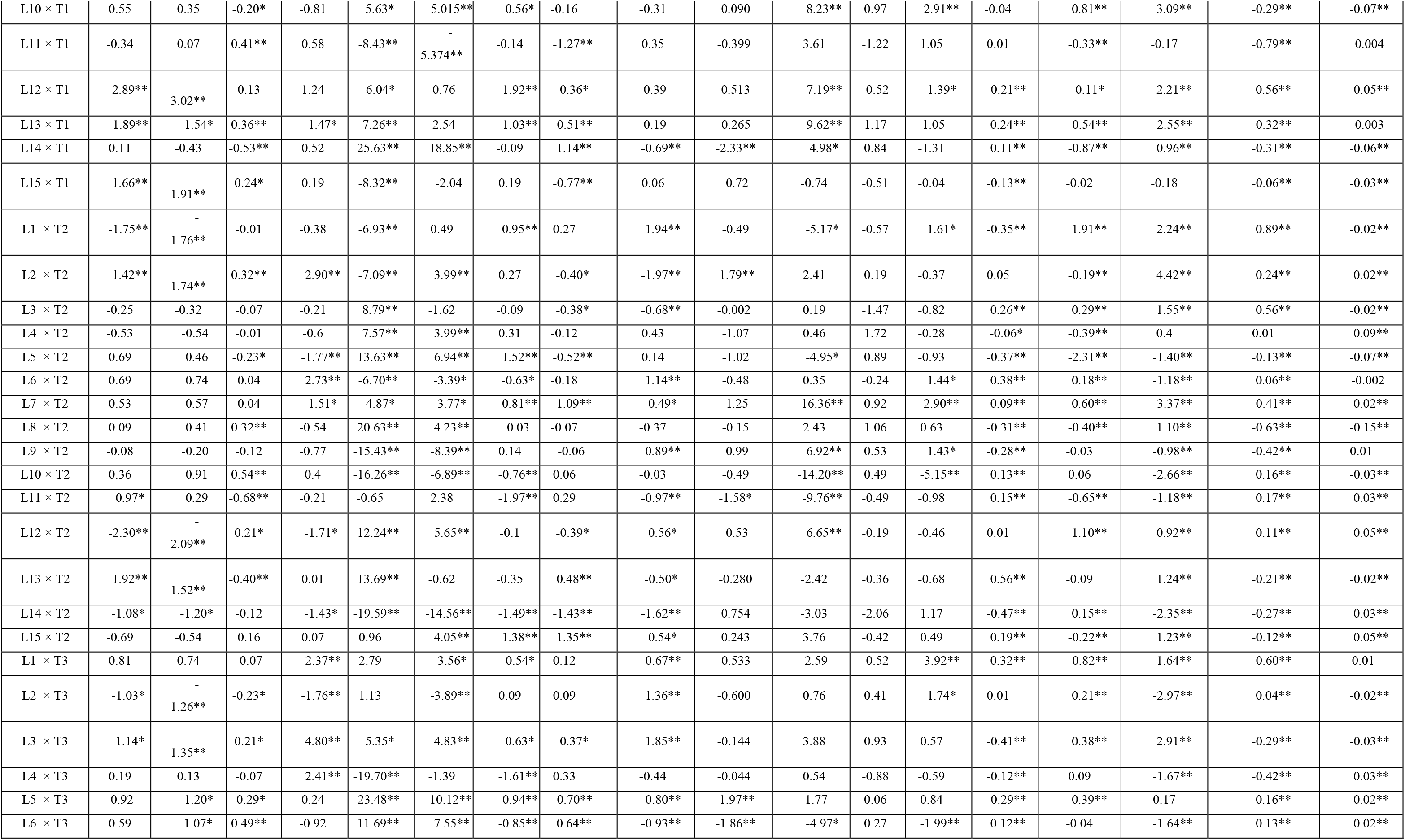
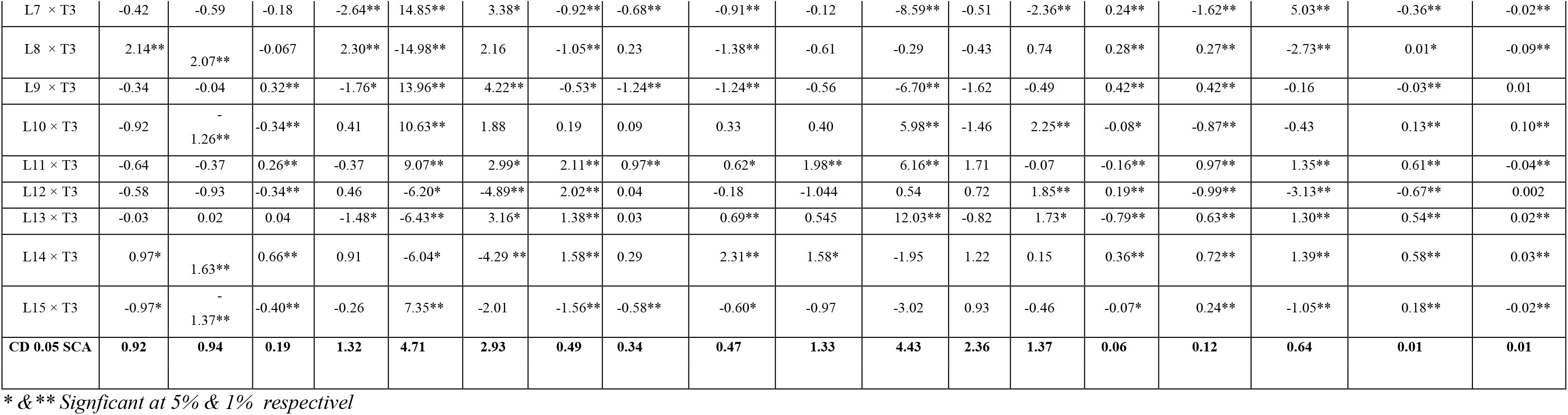
The specific combining abilities effects (SCA) for phenological, morphological grain yield and quality traits in QPM hybrids over the location.

Crosses L3 × T1, L7 × T3, L1 × T3, L4 × T1, L6 × T1, L5 × T2, L8 × T1, L2 × T3, L9 × T3, L12 × T2, L13 × T3, L14 × T2 displayed significant and negative SCA effects for days to 75 percent brown husk (Table 3). Positive and considerable GCA effects were exhibited by L14, L13, L15,T3 and L3 showing poor combiners for the same trait. Crosses L14 × T2 was showed highest negative and significant SCA effects followed by L5 × T3, L9 × T2, L7 × T1, L10 × T2 for ear height. Thirteen crosses namely L11 × T3, L12 × T3, L14 × T3, L5 × T2, L6 × T1, L15 × T2, L13 × T3, L4 × T1, L8 × T1, L1 × T2, L7 × T2, L3 × T3 and L10 × T1 showed positive and significant SCA for ear length. The parent T3 involved in these crosses exhibited low GCA effects for ear height. Crosses L14 × T3, L2 × T2, L5 × T3, L11 × T3 and L6 × T1 showed positive and significant SCA effects for 100-grain weight. Whereas crosses L11 × T2, L6 × T3, L14 × T1 showed negative and significant SCA effects (Table 3). Eleven crosses *viz.,* L7 × T2, L13 × T3, L10 × T1, L1 × T1, L9 × T2, L5 × T1, L12 × T2, L11 × T3, L10 × T3, L14 × T1 and L6 × T1 had positive and significant SCA effects for grain yield. Significant and positive SCA effects were associated with a higher mean performance for grain yield in the progeny. Crosses L10 × T1, L7 × T2, L1 × T1, L10 × T3, L12 × T3, L2 × T3, L13 × T3, L1 × T2, L6 × T2, L9 × T2 recorded positive and significant SCA effects for harvest index.in a desirable direction. For oil content crosses L5 × T1, L13 × T2, L9 × T3, L6 × T2, L14 × T3, L1 × T3, L8 × T3, L3 × T2, L13 × T1, L7 × T3, L15 × T2, L12 × T3, L4 × T1, L3 × T1, L11 × T2, L10 × T2, L6 × T3, L14 × T1 and L7 × T2 showed positive and significant SCA effects (Table 3).

Crosses L5 × T1 was showed highest positive and significant SCA effects followed by L1 × T2 L12 × T2, L7 × T1 and L11 × T3 for protein content (Table 3). For starch content crosses L7 × T3 followed by L2 × T2, L10 × T1, L3 × T3, L6 × T1 and L1 × T2 were exhibited high positive and significant SCA effects (Table 3). Crosses L1 × T2 showed highest positive and significant SCA effects followed by L7 × T1, L8 × T1, L11 × T3, L14 × T3 and L12 × T1 for lysine content. The highest positive and significant SCA effects was observed for tryptophan content among crosses L13 × T1 followed by L8 × T1, L10 × T3, L4 × T2, L3 × T1, L12 × T2 and L15 × T2 (Table 3).

### Heterosis Estimates

Estimates of heterosis of five best hybrids for grain yield, yield-related traits and biochemical traits are presented in Table 4. Heterosis over the mid parent crosses L13 × T1 showed highest negative and significant followed by L6 × T1, L12 × T2, L14 × T2 and L2 × T1 for days to 50 per cent tasselling while crosses L13 × T1, L6 × T1, L12 × T2, L14 × T2 and L2 × T1 over better parent were exhibited for the same traits (Table 4). A perusal estimates of mid parent (MP) heterosis for plant height revealed that only one hybrid L10 × T2 exhibited negative significant (Table 4). Only two L10 × T2 and L9 × T3 hybrid exhibited negative significant heterobeltiosis for the same traits (Table 4). The estimates positive significant heterobeltiosis was observed in L2 × T2, L5 × T3, L11 × T3, L4 × T3 and L14 × T3 on the pooled basis with range varied from 25.19 (L14 × T3) to 32.98% (L2 × T2) for this trait (Table 4).

**Table 4.** Top five hybrids selected on the basis of heterosis for phenological, morphological grain yield and quality traits in QPM hybrids over the location

| DFT | DFS | ASI | DSBH | PH | EH | EL | EG | GR | GWT | GY | SP | HI | OC | PC | SC | LC | TP |
| --- | --- | --- | --- | --- | --- | --- | --- | --- | --- | --- | --- | --- | --- | --- | --- | --- | --- |
| Heterosis (Het) |  |  |  |  |  |  |  |  |  |  |  |  |  |  |  |  |  |
| L13 × T1<br>-8.56** | L6 × T1<br>-9.31** | L6 × T1<br>-42.86** | L14 × T2<br>-4.57** | L10 × T2<br>-5.53* | L7 × T1<br>8.69** | L12 × T3<br>49.41** | L7 × T2<br>29.20** | L8 × T1<br>47.80** | L11 × T3<br>36.93** | L11 × T1<br>111.75** | L12 × T3<br>13.16** | L11 × T1<br>66.38** | L5 × T1<br>46.85** | L1 × T2<br>52.99** | L14 × T1<br>7.13** | L14 × T3<br>85.01** | L8 × T1<br>28.34** |
| L6 × T1<br>-7.84** | L8 × T1<br>-7.81** | L14 × T1<br>-35.71** | L4 × T2<br>-4.18** | L9 × T3<br>5.80* | L9 × T2<br>9.66** | L14 × T3<br>44.32** | L14 × T1<br>27.09** | L11 × T3<br>42.86** | L12 × T2<br>36.86** | L7 × T2<br>110.23** | L12 × T1<br>13.02** | L7 × T2<br>62.89** | L11 × T2<br>20.32** | L5 × T1<br>35.66** | L9 × T1<br>5.87** | L11 × T3<br>53.67** | L3 × T1<br>23.97** |
| L12 × T2<br>-7.20** | L13 × T1<br>-7.35** | L3 × T1<br>-24.14** | L2 × T1<br>-3.54** | L3 × T2<br>7.46** | L4 × T1<br>10.90** | L11 × T3<br>42.88** | L9 × T1<br>25.57** | L11 × T1<br>41.59** | L3 × T2<br>35.77** | L5 × T1<br>109.13** | L11 × T3<br>10.84** | L12 × T3<br>59.97** | L11 × T1<br>18.23** | L11 × T3<br>30.62** | L14 × T3<br>2.69** | L12 × T1<br>39.08** | L5 × T1<br>23.85** |
| L14 × T2<br>-6.46** | L3 × T1<br>-7.1** | L9 × T1<br>-24.14** | L3 × T1<br>-3.54** | L8 × T2<br>8.41** | L14 × T2<br>11.27** | L1 × T2<br>41.24** | L13 × T2<br>23.69** | L14 × T3<br>39.79** | L3 × T3<br>34.51** | L12 × T2<br>108.17** | L12 × T2<br>10.32** | L10 × T1<br>59.89** | L10 × T2<br>18.05** | L14 × T3<br>29.86** | L1 × T2<br>1.95** | L4 × T1<br>36.02** | L4 × T2<br>22.41** |
| L2 × T1<br>-6.30** | L2 × T1<br>-6.79** | L8 × T1<br>-21.43** | L4 × T1<br>-3.11** | L15 × T2<br>15.32** | L1 × T3<br>15.39** | L13 × T3<br>38.19** | L14 × T3<br>20.10** | L3 × T3<br>37.37** | L5 × T3<br>34.45** | L10 × T1<br>105.01** | L14 × T1<br>8.59** | L4 × T1<br>58.27** | L13 × T2<br>17.65** | L5 × T3<br>28.9** | L7 × T3 -<br>0.15 | L7 × T1<br>33.62** | L11 × T2<br>20.25** |
| CD at 0.051.13 | 1.15 | 0.24 | 1.62 | 5.76 | 3.59 | 0.60 | 0.42 | 0.58 | 1.63 | 5.42 | 2.90 | 1.68 | 0.07 | 0.13 | 0.79 | 0.01 | 0.01 |
| CD at 0.01 1.49 | 1.52 | 0.32 | 2.14 | 7.60 | 4.74 | 0.79 | 0.55 | 0.76 | 2.15 | 7.15 | 3.82 | 2.22 | 0.09 | 0.17 | 1.04 | 0.01 | 0.01 |
| Heterobeltiosis (Hetb) |  |  |  |  |  |  |  |  |  |  |  |  |  |  |  |  |  |
| L13 × T1<br>-14.08** | L6 × T1<br>-12.21** | L6 × T1 -<br>50** | L4 × T2<br>-8.87** | L10 × T2<br>-11.36** | L4 × T1<br>-4.79** | L12 × T3<br>40.35** | L7 × T2<br>24.01** | L8 × T1<br>40 | L2 × T2<br>32.98** | L14 × T1<br>84.39** | L12 × T1<br>8.82** | L11 × T1<br>65.96** | L5 × T1<br>44.80** | L1 × T2<br>48.83** | L9 × T1<br>1.99** | L14 × T3<br>32.37** | L3 × T1<br>22.43** |
| L6 × T1<br>-11.45** | L13 × T1<br>-12.01** | L14 × T1<br>-43.75** | L4 × T1<br>-7.95** | L9 × T3<br>-2.50 | L7 × T1<br>1.59 | L11 × T3<br>36.93** | L9 × T1<br>21.43** | L11 × T1<br>38.51** | L5 × T3<br>30.52** | L5 × T1<br>81.97** | L12 × T3<br>7.71** | L10 × T1<br>56.44** | L11 × T2<br>19.79** | L11 × T3<br>29.65** | L14 × T1<br>1.37** | L1 × T2<br>19.93** | L5 × T1<br>21.62** |
| L12 × T2<br>-11.18** | L12 × T2<br>-10.69** | L3 × T1 -<br>31.25** | L2 × T1<br>-7.37** | L3 × T2<br>1.40 | L9 × T2<br>1.73 | L1 × T2<br>34.42** | L14 × T1<br>21.03** | L14 × T3<br>38.34** | L11 × T3<br>27.57** | L10 × T1<br>76.11** | L14 × T1<br>7.65** | L4 × T1<br>56.16** | L10 × T2<br>16.96** | L5 × T3<br>27.39** | L7 × T3 -<br>0.72 | L11 × T3<br>2.22** | L11 × T2<br>19.5** |
| L14 × T2<br>-10.85** | L14 × T2<br>-10.48** | L8 × T1 -<br>31.25** | L3 × T1<br>-7.37** | L9 × T1<br>4.50 | L10 × T2<br>3.47 | L14 × T3<br>34.29** | L15 × T2<br>17.27** | L3 × T3<br>36.65** | L4 × T3<br>25.29** | L11 × T1<br>69.78** | L14 × T3<br>7.28** | L7 × T1<br>54.45** | L13 × T2<br>16.95** | L12 × T2<br>25.91** | L6 × T1 -<br>1.83** | L14 × T2<br>-5.36** | L4 × T2<br>18.92** |
| L2 × T1<br>-9.73** | L2 × T1<br>-9.38** | L9 × T1 -<br>31.25** | L14 × T2<br>-7.34** | L8 × T2<br>6.82* | L4 × T3<br>4.57* | L10 × T1<br>31.94** | L13 × T2<br>16.05** | L11 × T3<br>29.63** | L14 × T3<br>25.19** | L4 × T1<br>69.65** | L13 × T1<br>7.02** | L7 × T2<br>54.45** | L6 × T2<br>15.41** | L7 × T2<br>25.47** | L8 × T1 -<br>2.18** | L12 × T2<br>-5.55** | L15 × T2<br>17.38** |
| CD at 0.05, 1.30 | 1.33 | 0.28 | 1.87 | 6.65 | 4.15 | 0.69 | 0.48 | 0.67 | 1.88 | 6.26 | 3.34 | 1.94 | 0.08 | 0.15 | 0.91 | 0.01 | 0.01 |
| CD at 0.01, 1.72 | 1.76 | 0.37 | 2.47 | 8.78 | 5.47 | 0.91 | 0.63 | 0.88 | 2.48 | 8.26 | 4.41 | 2.56 | 0.11 | 0.20 | 1.20 | 0.02 | 0.02 |

\* & \*\* Significant at 5% & 1% respectivel

Significant and positive mid parent heterosis were manifested by L11 × T1, L7 × T2, L5 × T1, L12 × T2 and L10 × T1 varied from 105.01% (L10 ×T1) to 111.75% (L11 × T1) for grain yield per plant. Increase heterobeltiosis by significant positive heterosis was exhibited L14 × T1 with a magnitude of range varied from 69.65% (L4 ×T1) to 84.39% (L14 × T1) for the grain yield (Table 4). The heterosis over mid parent in positive direction ranged from 17.65% (L13 × T2) to 46.85% (L5 × T1) and heterobeltiosis varied from 15.41% (L6 × T2) to 44.80% (L5 × T1) for oil content. Heterosis over mid parent in positive direction ranged from 28.90 (L5 × T3) to 52.99% (L1 × T2) and heterosis over parents varied from 25.47 (L7 × T2) to 48.83% (L1 × T2) for protein content. The perusal of estimates of significant positive relative heterosis for this starch content was recorded in hybrids namely L14 × T1, L9 × T1, L14 × T3, L1 × T2 and L7 × T3 (Table 4). The highest estimates of significant standard heterosis in a positive direction were exhibited by hybrids L1 × T2, L14 × T3, L12 × T1, L7 × T1 and L4 × T1 for lysine content. The relative heterosis in positive direction ranged from 20.25 (L11 × T2) to 28.34% (L8 × T1) for tryptophan content and significant positive heterobeltiosis with range varied from 17.38 (L15 × T2) to 22.43% (L3 × T1) for this trait (Table 4).

### Heterotic Groupings

The dendrogram constructed based on the GCA effects revealed three heterotic groups across location (Figure 2). Group, I consisted of inbred line L5, L15, L10 and L13; group II comprises, L14, L12, L11 and T3; group III consist of L1, L7 and L9 while, L4, T1, T2, L3, L2, L6 and L8 constituted group IV (Figure 2). According to Pswarayi and Vivek [22], higher GCA effects, are ideal for the classification into a heterotic group and per se grain yield. Based on these criteria, L2, L6, L4, L8, T1and L3 were the highest yielding parents across location (Table 2), was placed in the fourth heterotic group (Figure 2). Also, Inbred line L14 and L12 recorded high grain yield, had the highest positive GCA effects for grain yield, was classified into the second heterotic group and was therefore identified as the best tester for heterotic group second whereas inbred line L15 and L5 were identified as the best tester in the heterotic first group (Figure 2).

**Figure 2.**
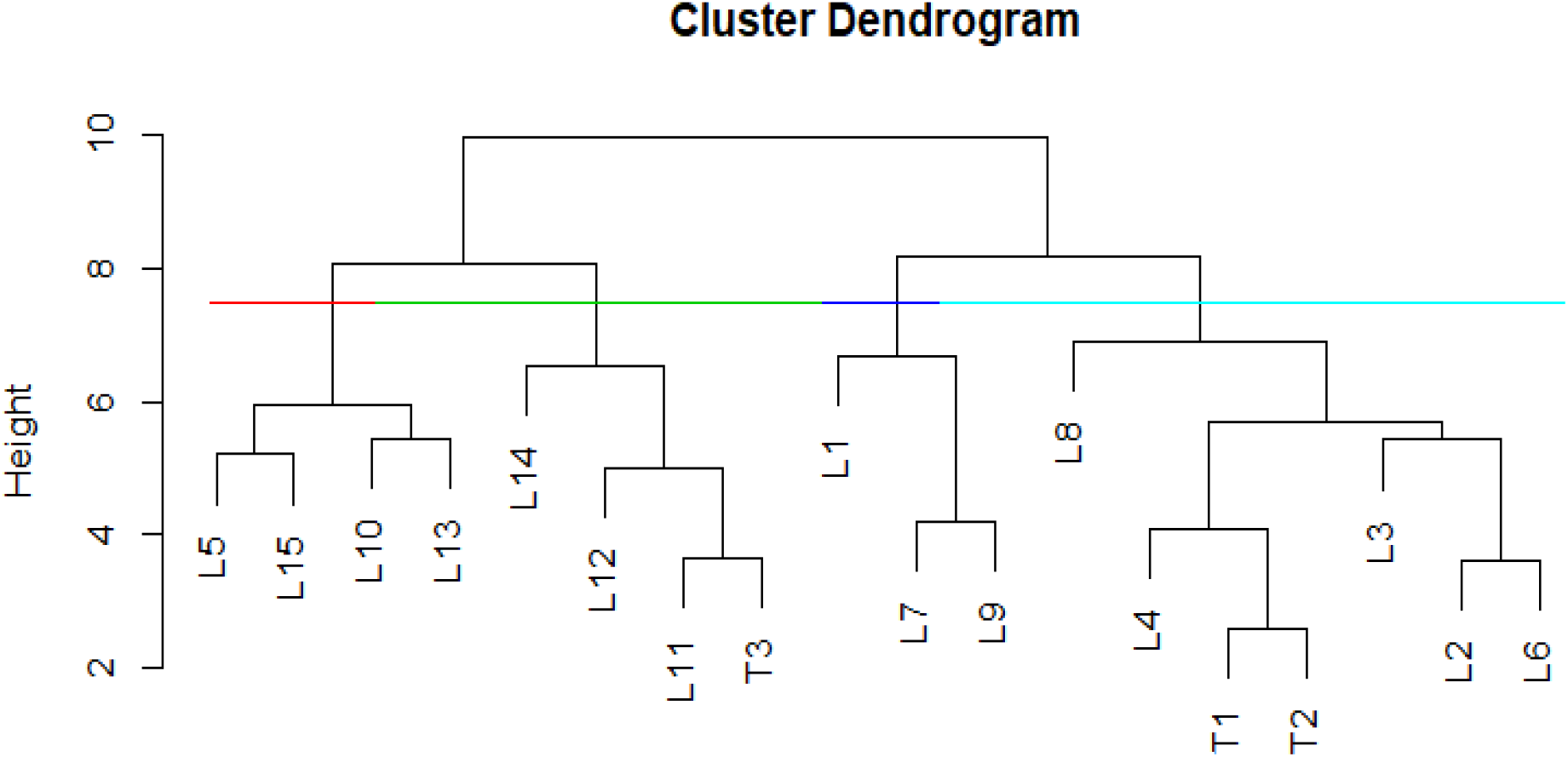
Dendrogram of 15 inbred lines and 3 testers constructed from GCA effects of grain yield and multiple traits using Ward’s minimum variance cluster analysis across environments

### Components of genetic variance

SCA variance was larger than average GCA by the average σ^2^ GCA/σ^2^ SCA ratio being less than one for the all traits (Table 5). The predictability ratios more than 0.50 for only days to 75 percent brown husk indicating additive gene action. Whereas all traits except days to 75 percent brown husk were recorded less than 0.50, pointing out for the non-additive gene action (Table-5). Maximum genetic advance exhibited by plant height (11.00%) followed by ear height (5.60%), days to 75 percent brown husk and grain yield (2.40%). Hybrids showed maximum contribution (%) over the line and tester for all the traits (Table 5).

**Table 5.**
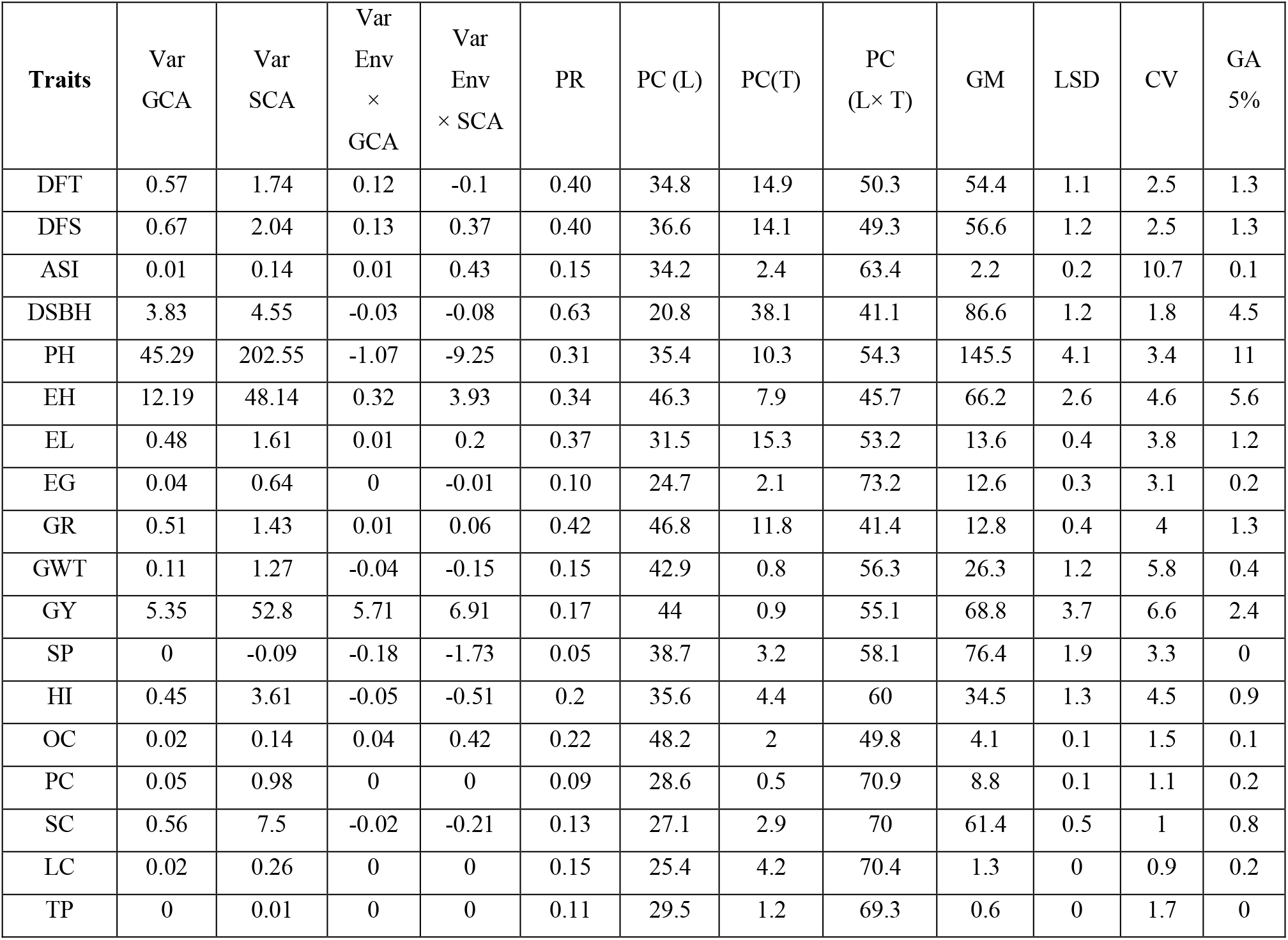
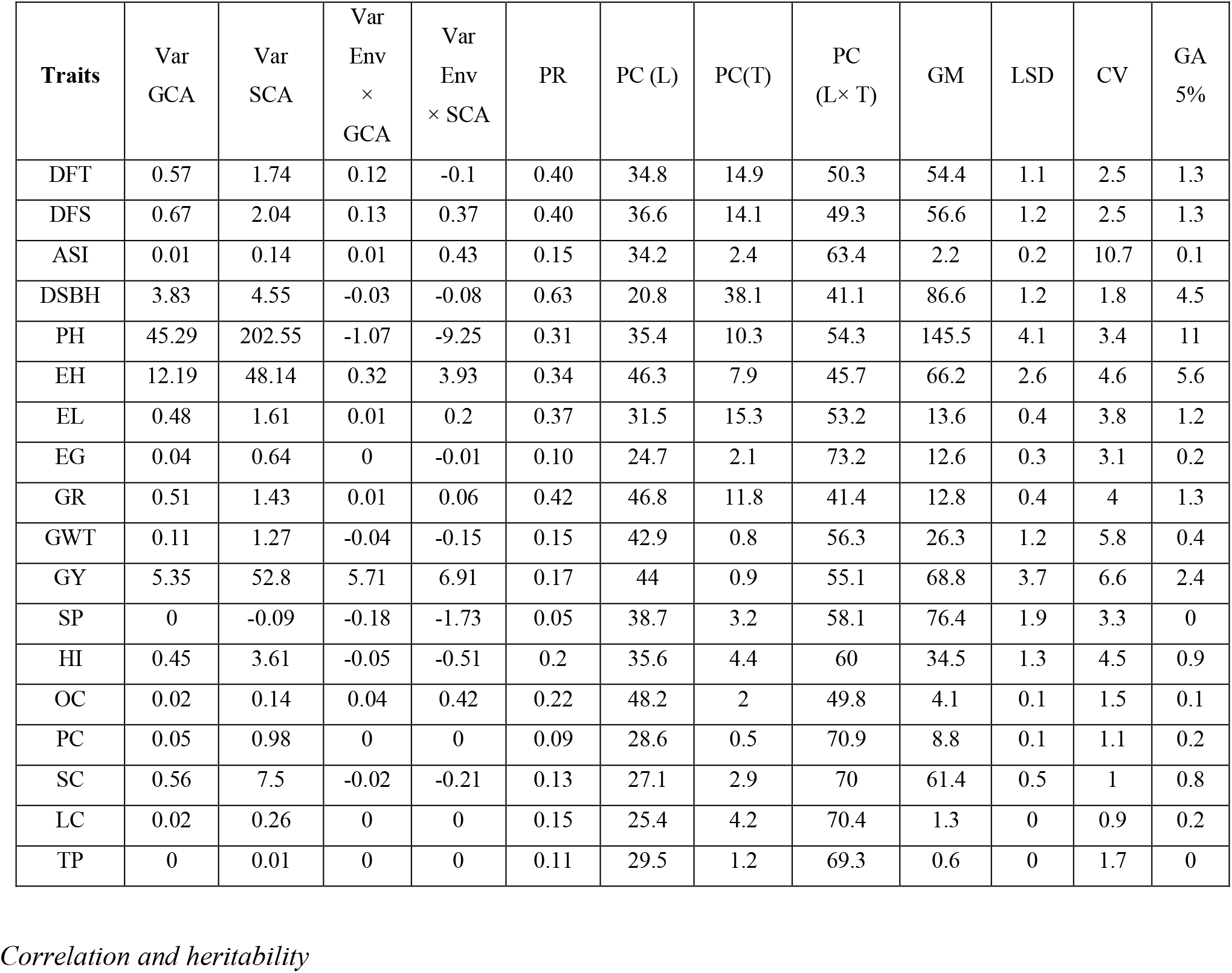
Components of genetic variance, genetic advance, the contribution of line, tester and their hybrids in QPM.

| Traits | Var<br>GCA | Var<br>SCA | Var<br>Env<br>×<br>GCA | Var<br>Env<br>× SCA | PR | PC (L) | PC(T) | PC<br>(L× T) | GM | LSD | CV | GA<br>5% |
| --- | --- | --- | --- | --- | --- | --- | --- | --- | --- | --- | --- | --- |
| DFT | 0.57 | 1.74 | 0.12 | -0.1 | 0.40 | 34.8 | 14.9 | 50.3 | 54.4 | 1.1 | 2.5 | 1.3 |
| DFS | 0.67 | 2.04 | 0.13 | 0.37 | 0.40 | 36.6 | 14.1 | 49.3 | 56.6 | 1.2 | 2.5 | 1.3 |
| ASI | 0.01 | 0.14 | 0.01 | 0.43 | 0.15 | 34.2 | 2.4 | 63.4 | 2.2 | 0.2 | 10.7 | 0.1 |
| DSBH | 3.83 | 4.55 | -0.03 | -0.08 | 0.63 | 20.8 | 38.1 | 41.1 | 86.6 | 1.2 | 1.8 | 4.5 |
| PH | 45.29 | 202.55 | -1.07 | -9.25 | 0.31 | 35.4 | 10.3 | 54.3 | 145.5 | 4.1 | 3.4 | 11 |
| EH | 12.19 | 48.14 | 0.32 | 3.93 | 0.34 | 46.3 | 7.9 | 45.7 | 66.2 | 2.6 | 4.6 | 5.6 |
| EL | 0.48 | 1.61 | 0.01 | 0.2 | 0.37 | 31.5 | 15.3 | 53.2 | 13.6 | 0.4 | 3.8 | 1.2 |
| EG | 0.04 | 0.64 | 0 | -0.01 | 0.10 | 24.7 | 2.1 | 73.2 | 12.6 | 0.3 | 3.1 | 0.2 |
| GR | 0.51 | 1.43 | 0.01 | 0.06 | 0.42 | 46.8 | 11.8 | 41.4 | 12.8 | 0.4 | 4 | 1.3 |
| GWT | 0.11 | 1.27 | -0.04 | -0.15 | 0.15 | 42.9 | 0.8 | 56.3 | 26.3 | 1.2 | 5.8 | 0.4 |
| GY | 5.35 | 52.8 | 5.71 | 6.91 | 0.17 | 44 | 0.9 | 55.1 | 68.8 | 3.7 | 6.6 | 2.4 |
| SP | 0 | -0.09 | -0.18 | -1.73 | 0.05 | 38.7 | 3.2 | 58.1 | 76.4 | 1.9 | 3.3 | 0 |
| HI | 0.45 | 3.61 | -0.05 | -0.51 | 0.2 | 35.6 | 4.4 | 60 | 34.5 | 1.3 | 4.5 | 0.9 |
| OC | 0.02 | 0.14 | 0.04 | 0.42 | 0.22 | 48.2 | 2 | 49.8 | 4.1 | 0.1 | 1.5 | 0.1 |
| PC | 0.05 | 0.98 | 0 | 0 | 0.09 | 28.6 | 0.5 | 70.9 | 8.8 | 0.1 | 1.1 | 0.2 |
| SC | 0.56 | 7.5 | -0.02 | -0.21 | 0.13 | 27.1 | 2.9 | 70 | 61.4 | 0.5 | 1 | 0.8 |
| LC | 0.02 | 0.26 | 0 | 0 | 0.15 | 25.4 | 4.2 | 70.4 | 1.3 | 0 | 0.9 | 0.2 |
| TP | 0 | 0.01 | 0 | 0 | 0.11 | 29.5 | 1.2 | 69.3 | 0.6 | 0 | 1.7 | 0 |

### Correlation and heritability

The results of the correlation analysis for grain yield, its contributing and biochemical traits are presented in Table 6. Oil content and days to days to 75 per cent brown husk were showed significant and positive association with grain yield. F1 means showed highly significant correlation with r(F1,Het), r(F1,hebt), for all the traits (Table 6).Correlation between F1mean and SCA r(F1, SCA) also exhibited a significant correlation for only lysine content. Correlation between SCA and Het (rSCA,Het) showed negative and significant for only starch and lysine content (Table 6). Heritability (bs) ranged from 82.80% (Shelling percentage) to 99.99% (Lysine content) and all traits exhibited more than 90% heritability except shelling percentage (Table 5). Lysine content showed highest moderate heritability(ns) followed by starch content, 100-grain weight, grain yield, tryptophan content and ear girth (Table 6).

**Table 6.** Average mid-parent heterosis correlation among F1, heterosis (Het.), heterobeltiosis (Heb), general and specific combining ability (SCA) effects for 45 QPM single crosses.

| Traits | r (F1, Het) | r (F1,Heb) | r (F1, SCA) | r (F1, GCA) | r (Het, SCA) | AH (%) | F test |
| --- | --- | --- | --- | --- | --- | --- | --- |
| DFT | 0.74** | 0.53** | 0.19 | 0.34* | 0.22 | -1.50 | ** |
| DFS | 0.79** | 0.61** | 0.23 | 0.39** | 0.21 | -1.22 | ** |
| ASI | 0.87** | 0.81** | 0.11 | 0.12 | 0.06 | 6.84 | ** |
| DSBH | 0.88** | 0.73** | 0.20 | 0.31* | 0.20 | 1.73 | ** |
| PH | 0.79** | 0.74** | 0.11 | 0.03 | 0.12 | 30.09 | ** |
| EH | 0.54** | 0.62** | 0.14 | 0.18 | 0.16 | 34.88 | ** |
| EL | 0.89** | 0.83** | 0.16 | 0.02 | 0.19 | 23.42 | ** |
| EG | 0.80** | 0.84** | -0.04 | 0.26 | -0.12 | 11.99 | ** |
| GR | 0.86** | 0.86** | 0.12 | -0.08 | 0.14 | 15.18 | ** |
| GWT | 0.23 | 0.32* | 0.22 | 0.13 | -0.04 | 19.97 | ** |
| GY | 0.62** | 0.64** | 0.16 | -0.20 | 0.10 | 64.51 | ** |
| SP | 0.74** | 0.71** | 0.12 | 0.11 | 0.05 | 4.44 | ** |
| HI | 0.57** | 0.41** | 0.04 | 0.00 | -0.05 | 45.86 | ** |
| OC | 0.86** | 0.81** | 0.16 | 0.12 | 0.16 | 4.19 | ** |
| PC | 0.86** | 0.76** | -0.14 | -0.43** | -0.14 | 9.78 | ** |
| SC | 0.68** | 0.75** | -0.13 | -0.11 | -0.25* | -5.07 | ** |
| LC | 0.89** | 0.94** | -0.28* | -0.12 | -0.27* | -13.81 | ** |
| TP | 0.61** | 0.33* | 0.11 | -0.14 | 0.19 | 4.95 | ** |

*DFT: Days to 50 per cent tasselling, DFS:Days to 50 per cent silking, ASI: Anthesis silking interval, PH: Plant height (cm), EH: Ear height (cm), EL: Ear length (cm), EG: Ear girth (cm), GR: Number of grain rows per ear , GWT: 100-grain weight (g), GY: Grain yield per plant (g), SP: Shelling percentage (%),HI: Harvest index (%) , OC: Oil content (%),PC: Protein content (%),SC: Protein content (%),LC: Lysine content (%),TP: Tryptophan content (%)*
*AH: Average Heterosis , PR: Predictability ratios, GA: Genetic Advance, LSD: Least Significant Differences, GM= General mean, CV: Coefficient of Variation; \* & \*\* Significant at 5% & 1% respectively*

## Discussion

Development of maize genotypes with high yield potential would enhance production and productivity. Assessment of heterosis and combining ability is useful for developing superior hybrids for improving yield and quality traits [23–25]. The current study improving grain yield and nutritional value of quality protein maize to identify suitable hybrids for further selection and breeding. Many researchers have also reported significant variations among genotypes, environments and their interactions for yield,yield-related traits and quality traits in maize [23–27]. Line by location, tester by location, and line × tester by location line tester and line × tester main effects were significant for most traits. Highly significant line, tester and line × tester effects suggested both additive and non-additive effects of the environment on trait expression. Population development is dependent on crosses using complementary genotypes possessing desirable yield, yield component traits and quality traits. In line with the present study, maize progenies were previously reported to flower and mature early than parental genotypes due to additive gene effects in the inheritance of these traits [25,28]. Parents L7, L9, L11, L8, T2, and L2 good general combiner for plant height and their crosses L5 × T3 followed by L4 × T3, L14 × T2, L10 × T2, L9 × T2, L8 × T3 and L3 × T1 had highest negative significant SCA effects for plant height has been reported previously [23,25,28]. Crosses L14 × T3, L2 × T2, L5 × T3, L11 × T3 and L6 × T1 showed positive and significant SCA effects for the 100-grain weight which were greater than other crosses and some parental genotypes (Table 3).

These suggested parental genotypes contributed to enhanced 100-grain weight in the progenies. Crosses produced higher grain yield, which varied from 62.80g/plant (L9 × T3) to 92.12g/plant (L7 × T2) which was higher than parental genotypes. Similar to the present findings, many scientists reported higher grain yield among maize hybrids compared to parental genotypes [25,28-30]. The parental genotypes, suggesting heterosis for these traits in agreement with the previous reports [31]. Significant variation for heterosis and heterobeltiosis was observed for the studied agronomic and quality traits among crosses indicating genetic divergence to identify promising crosses for breeding. Positive and highly significant estimates of relative heterosis were manifested by L14 × T3, L11 × T3, L12 × T1, L4 × T1 and L7 × T1 for lysine content. Similar finding reported by [29,30]. Grain yield also exhibited high correlation with plant height, ear height, ear height, ear girth, number of grain rows per ear, 100-grain weight, shelling percentage and harvest index. This results conformity with [32–34]. Furthermore, a correlation between F1 mean and SCA r(F1, SCA) also exhibited significant correlation for only lysine content. These results were in agreement with [35–37] that emphasized the higher efficiency of SCAs for predicting the grain yield of QPM maize hybrids. Heritability(bs) all traits exhibited more than 90% heritability except shelling percentage. Whereas heritability (ns) lysine content showed highest moderate heritability(ns) followed by starch content, 100-grain weight, grain yield, tryptophan content and ear girth similar finding had reported by [33].

Predictability ratios more than 0.50 value for only days to 75 percent brown husk indicating additive gene action. Whereas all traits except days to 75 percent brown husk were recorded less than 0.50 value, indicating the predominance of non-additive gene action for these characters. The studies in the past confirmed similar results [30,31]. Similarly, variance due to GCA interaction with an environment lower than variance due to SCA interaction with the environment for all the trait. Similar finding proposed by [25,31]. Maximum genetic advance exhibited by plant height followed by ear height, days to 75 percent brown husk and grain yield. Hybrid\s showed maximum contribution over the line and tester for all the traits. Similar finding reported by [38].

## Conclusions

The present study Improving grain yield and nutritional value of quality protein maize (*Zea mays* L.) by appropriate hybrid selection. For further selection and breeding. Maize families such., L7 × T2, L13 × T3, L10 × T1, L1 × T1, L9 × T2, L5 × T1, L12 × T2, L11 × T3, L10 × T3, L14 × T1 and L6 × T1 were identified with suitable yield and component traits. Crosses L5 × T1, L3 × T1, L12 × T2, L7 × T3, L1 × T2 and L13 × T1 were identified for quality traits. These are useful maize populations for direct production.

## Acknowledgement

This study was supported by the All India Co-ordinated Maize Improvement Project, MPUAT, Udaipur. The authors thank principal investigator All India Co-ordinated Maize Improvement Project, MPUAT, Udaipur for providing experimental material, valuable help and support. The authors also thank UGC to provide the Rajiv Gandhi National Fellowship (F1•17.1/2014•15/RGNF•2014•15•SC•UTT•57779/SA•III/Website) during research work for financial support.

